# Federated queries of clinical data repositories: balancing accuracy and privacy

**DOI:** 10.1101/841072

**Authors:** Yun William Yu, Griffin M Weber

## Abstract

Researchers use large federated clinical data networks that connect dozens of healthcare organizations to access data on millions of patients. However, because patients often receive care from multiple sites in the network, queries frequently double-count patients. Using the probabilistic streaming algorithm HyperLogLog and adding obfuscation, we developed a scalable method for estimating the number of distinct lives that match a query, which balances accuracy and privacy in a “tunable” way.

## I. INTRODUCTION

Widespread adoption of electronic health records has generated vast amounts of data, which are increasingly being used in clinical research, epidemiological studies, and public health applications [Jensen, et al., 2012]. Data from multiple healthcare organizations are often needed in order to increase statistical power or to access diverse patient populations and geographic regions. However, HIPAA and other patient privacy laws generally prevent data from different sites from being combined into a central repository for analysis. Therefore, several federated clinical data research networks have been developed that broadcast queries to multiple sites, run analyses locally, and then combine the results. Two of the largest in the United States are the PCORNet network for patient-centered outcomes research [Fleurence 2014] and the NIH-funded Accrual for Clinical Trials (ACT) network [Weber 2009, McMurry 2013, Visweswaran 2018], both of which connect dozens of healthcare organizations across the country and include health data on nearly 100 million Americans.

Because patients often receive care at more than one clinical site, the data for a patient at any one site might not be complete, and the same information about a patient might be duplicated at different sites. This can lead to queries returning incorrect results. A similar situation arises when patients’ data are intentionally separated for technical reasons, such as when large amounts clinical data (e.g., diagnoses and medications) and genomic data are stored in different locations, and it is not feasible to merge them into a single database. In both cases, computation must be performed in a distributed fashion, but the challenge is that an individual patient’s data may be spread across multiple databases.

Various methods to addressing this problem have been described in the literature, but they have different tradeoffs in terms of accuracy, privacy, scalability and computational complexity. We group these into three broad categories (aggregate counts, hashed patient identifiers, and privacy-guaranteed methods) and introduce a new, better balanced approach. To compare these methods, we developed a generalizable benchmarking framework and software that simulates queries of large federated data networks.

### Aggregate counts

Federated queries in PCORNet and ACT ask sites to return the number of patients in their local databases who match some set of criteria, such as having both hypertension and diabetes. The networks present the user with the aggregate count from each site; and, no attempt is made to link patients across sites or de-duplicate records. This can lead to large overestimates of the number of distinct patients who match a query if the counts from each site are naively summed [Weber 2013]. To protect patient privacy, the networks mask small counts by displaying “<10 patients”. However, the results from multiple queries can be combined to reveal information about individual patients (see Methods for details). Sites participating in these networks are aware of this privacy risk, which they mitigate through institutional agreements that require sites to audit researchers’ queries and monitor their use of the network.

### Hashed Patient Identifiers

The most accurate and semi-secure way to de-duplicate the results in a federated query is for each site to return the full list of patients who match the query. Privacy is the main concern since data on every patient matching the query (potentially many millions of people) must be shared. Patient identifiers (e.g., name and date of birth) are typically encrypted using a one-way hash function, such as SHA-1 [Eastlake 2001]. The same patient at two sites will be hashed to the same value if the same hash function is used (and there are no inconsistencies in the underlying demographic data). Unfortunately, hash functions are vulnerable to “dictionary attacks”, where an adversary who knows the encryption method can simply generate a “rainbow table” of the hashes of every possible patient identifier (e.g., all 9-digit social security numbers) and then use this to re-identify the list of hash values returned by a site [Oechslin 2003].

### Privacy-guaranteed methods

Secure multi-party computation (MPC) and homomorphic encryption techniques enable true privacy guarantees in a federated network (see Methods), and have recently been introduced for distributed genome-wide association studies [Cho, 2018] and pharmacological collaboration [Hie & Cho, 2018]. The limitation of these algorithms is computational complexity. Protocols that securely determine the number of shared patients between two sites [Kolesnikov 2017, de Cristofaro 2012, Weber 2013, Swamidass 2015] are impractical for large networks since the number of pairwise and multi-way comparisons grows exponentially with the number of sites. Other approaches that avoid exponential comparison either require sharing gigabytes of data [Fenske 2017], making numerous rounds of back-and-forth communication [Dong 2017], or using trusted 3^rd^ parties [Yigzaw 2017]. These are also problematic because, as we have previously shown [Weber 2015], large federated clinical data networks are fragile, with multiple sites typically failing to respond even to aggregate count queries.

## II. RESULTS

In this paper we propose a new method based on the HyperLogLog (HLL) probabilistic sketching algorithm [Flajolet 2007]. Although HLL is widely used in many applications, such as internet search engines, to our knowledge it has not been applied to federated queries of health data. The basic idea behind HLL (and other k-minimum value sketches [Bar-Yossef 2002]) is that the minimum of a collection of random numbers between 0 and 1 is inversely proportional to how many numbers are present. For example, a single random number between 0 and 1 has expected value 0.5, but if we have 99 random numbers, the minimum has expected value 0.01. By using a hash function that maps patients to a random number between 0 and 1, we can estimate the number of patients who match a query by keeping track of just the minimum hash value. Note that the smallest of these values is also the minimum of all the patients hashes across all sites. If each site returns its minimal hash value, we can estimate the number of distinct patients in the whole network that match the query from the smallest value. While it may seem unintuitive that the network minimum hash is the same as the hash for one hospital, which hospital that minimum hash corresponds to changes when you use multiple hash functions, allowing the estimator to be accurate.

HLL improves the accuracy of this method by efficiently dividing the patients into k partitions and returning the minimum value of in each partition. HLL also returns the base-2 order of magnitude of the minimum values rather than the actual values, which greatly reduces the risk of re-identification from a dictionary attack. For k partitions, the relative error of HLL is approximately 1/sqrt(k). For example, asking sites to share a HLL sketch with only 100 values, the number of distinct patients can be estimated with a 10% relative error. Error can be reduced by increasing k. Although higher k increases the risk of re-identification, the risk is quantifiable and predictable, enabling networks to define policies that maximizes accuracy while reducing risk to an acceptable level. Using various obfuscation techniques, such as masking the sketches of certain sites (HLL+Mask), using different cryptographic “salt” for each query (HLL+Rehash), and scrambling the order of the values in the sketch (HLL+Shuffle), we can further reduce risk. Even if re-identification is possible, then information on only k patients is revealed, not every patient who matches the query.

To quantitatively measure privacy loss, we use an adapted m-anonymity model of privacy [Sweeney 2002; El Emam 2008] (see Methods: Privacy Risk), and run benchmarks for runtime, accuracy, and privacy-loss on shared hashed identifiers, sharing aggregate counts, and our proposed HLL approach. Existing privacy guaranteed methods do not scale well and are infeasible for running on large datasets, with either extremely high runtime or error, so we only compare our algorithm qualitatively to those three.

To do so, we developed software for generating simulated networks containing up to 100 sites and 100 million distinct patients and running queries using aggregate counts (*Count* and *Count+Mask*), shared hash values (*HashedIDs*), secure MPC, and HLL with and without obfuscation. HLL is implemented using *k*=2^1^=2 (*HLL1*), *k*=2^4^=16 (*HLL4*), *k*=2^7^=128 (*HLL7*), and *k*=2^15^=32,768 (*HLL15*). We provide all code in GitHub (https://github.com/yunwilliamyu/secure-distributed-union-cardinality).

Experimental details and theoretical proofs are provided in the Methods section, and raw results are furnished in the Supplement Tables 1-10. Here, Table 1 provides qualitative summary comparisons of the relative run times, accuracy, and privacy risk of the different methods. We use an adapted *m*-anonymity model of privacy, whereby the privacy risk is defined to be the number of revealed data points that correspond to fewer than 10 patients (Methods: Privacy risk score). HLL with obfuscation is significantly more accurate than aggregate counts, lower risk than sharing hash values of all matching patients, and more scalable than privacy guaranteeing algorithms. Figure 1 illustrates the accuracy-risk tradeoff between the algorithms quantitatively when each patient’s data are at two sites on average. Counts and hash values form the bounds of the graph, forcing networks to make extreme choices between accuracy and privacy. Variations of HLL fill in the remaining space, enabling networks to “tune” the method to achieve a more desirable balance for a given application.

**TABLE I.** Qualitative comparison of approaches to determining the number of distinct patients in a federated network who match a query. “Risk-Hub” is the risk to patient privacy if an adversary gains access to only the hub. “Risk-Hub+Site” is the risk if the adversary also gains access to one of the sites in the network. An asterisk (“*”) means obfuscation does not change the value. For example, Count and Count+MPC have the same error. “HLL7+MPC” means the value is the same as “HLL7+MPC” without shuffling. Blue, green, yellow and orange correspond to zero, (very) small, medium, and (very) large, respectively. “(-)” means the value gets smaller when more patients match a query, and “(+)” means the value gets larger.

| Method | Obfuscation | Approximation Error | Runtime Wait | Risk-Hub | Risk-Hub+Site |
| --- | --- | --- | --- | --- | --- |
| Count | None | large | very small | medium(-) | medium(-) |
|  | Mask | large | very small | zero | zero |
|  | MPC | * | medium | zero | zero |
| HLL7 | None | medium | small | medium(+) | medium(+) |
|  | Shuffle | * | * | small(+) | * |
|  | Rehash | * | medium(+) | zero | * |
|  | Mask | medium(+) | medium(-) | zero | zero |
|  | MPC | * | large | small(+) | small(+) |
|  | Shuffle+MPC | * | HLL7+MPC | very small(+) | HLL7+MPC |
| HLL15 | None | small | medium | large(+) | large(+) |
|  | Shuffle | * | * | small(+) | * |
|  | Rehash | * | medium(+) | zero | * |
|  | Mask | large(+) | medium(-) | zero | zero |
| HashedIDs | None | zero | medium(+) | very large(+) | very large(+) |
|  | Rehash | * | * | zero | * |

**Fig. 1.**
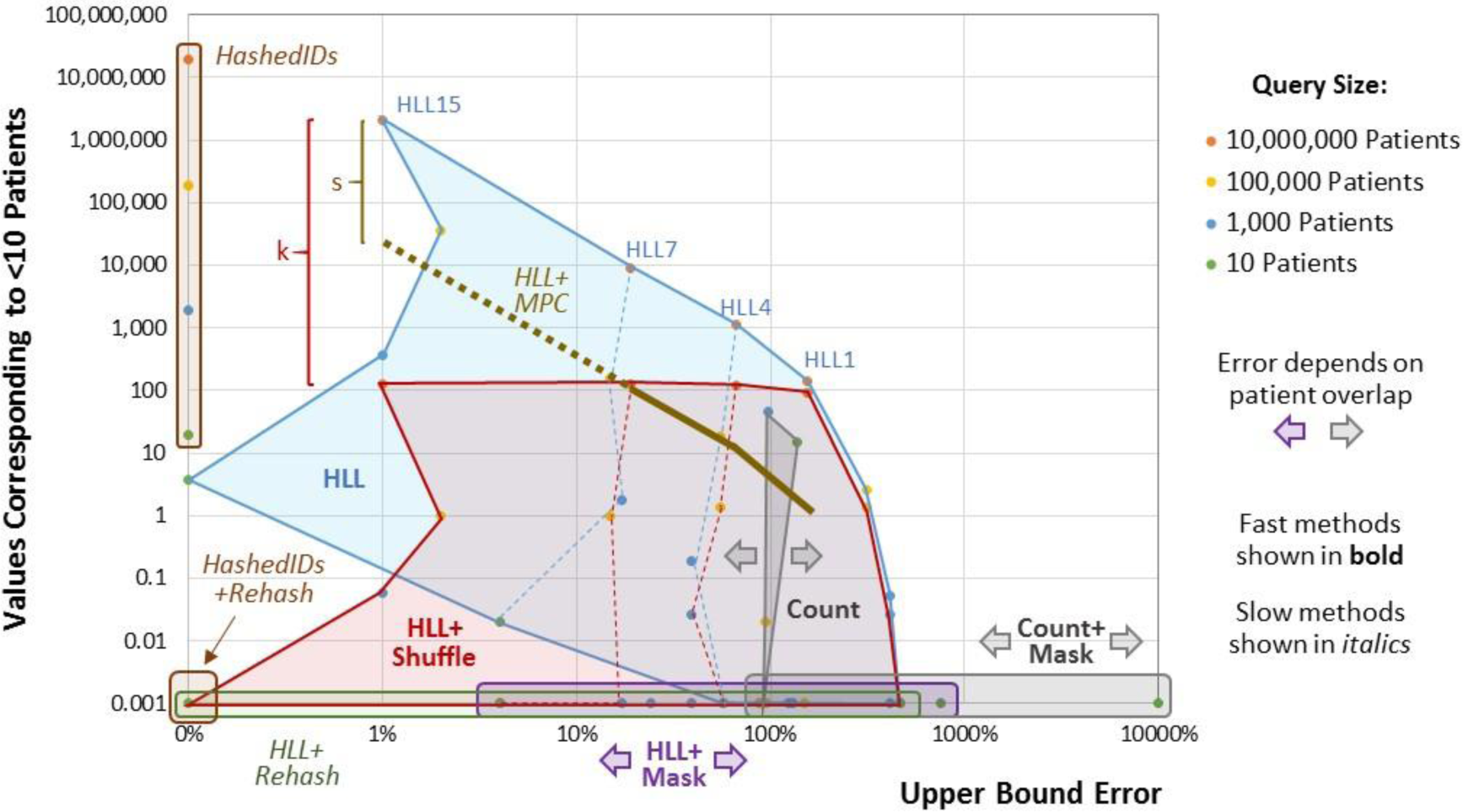
Quantitative graphical comparison of the query accuracy / privacy risk trade-off based on simulations of a network with 100 sites and 100 million patients. *HashedIDs* and *Count* bound the graph, while HLL-based methods enable a more balanced approach. (*HLL+MPC* is only shown for 10 million patients, and the values for *HLL7+MPC* and *HLL15+MPC* are theoretical rather than experimental.) *HLL+MPC* reduces the HLL risk by 1/*s*, where *s* is the number of sites in the network. *HLL+Shuffle* reduces the HLL risk by 1/*k*, where *k* is the number of values in the HLL sketch.

**Fig. 2.**
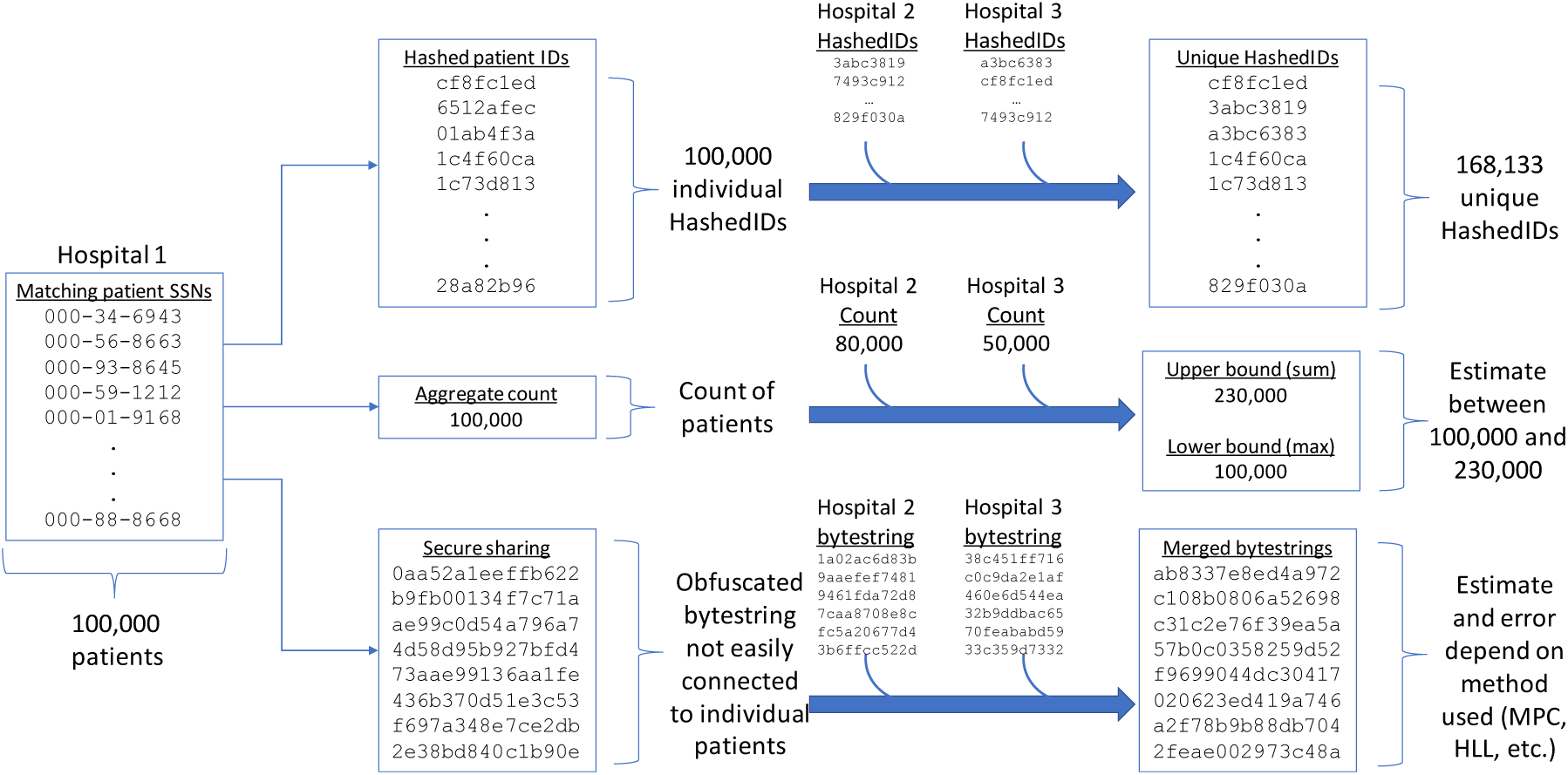
We classify methods for merging distributed queries into three groups: (1) sending full hashed patient identifiers, (2) sharing aggregate counts, and (3) generating obfuscated bytestrings that can be merged together. Sending hashedIDs gives an exact answer, but has a high privacy risk as each hash is associated with an individual patient. Sharing aggregate counts is relatively private, but has a high level of uncertainty and error in the answer. Obfuscated bytestring methods can range from probabilistic sketches to cryptographically secure MPC. They can provide privacy, but generally at the cost of computational complexity. In this paper, we provide much faster versions of obfuscated bytestring methods using HyperLogLog as a base.

**Fig. 3.**
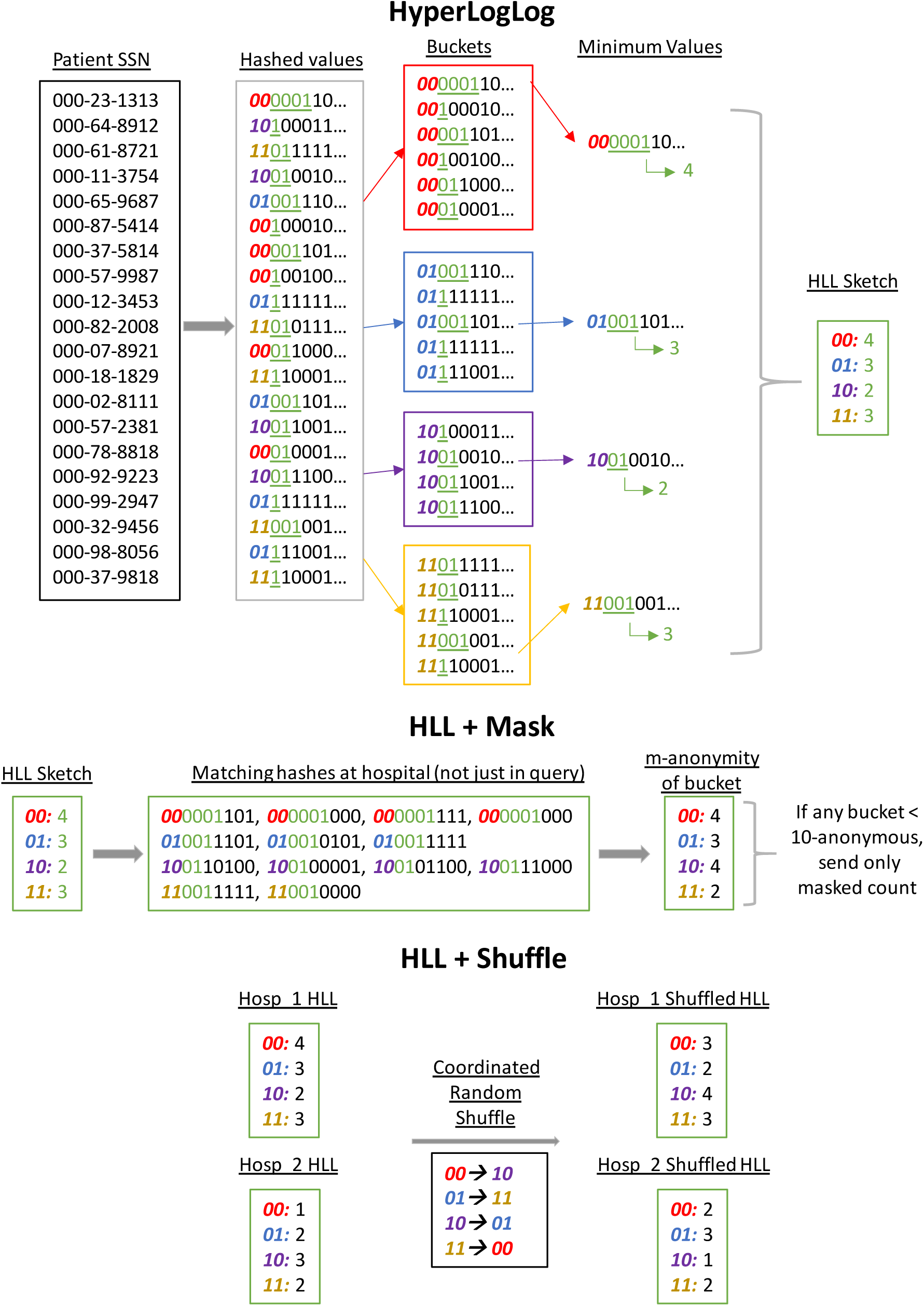
(top) HyperLogLog method. Given patient identifiers (e.g. SSN), we first hash them to a bitstring. The first several bits are used to bucket the values, and then within each bucket we store the position of the leading one indicator of the minimum value. (middle) HLL + Mask. We count the number of hashes that match the leading 1-indicator for each bucket; if that number is less than 10, the bucket is not 10-anonymous, so we do not send the HLL, but only a masked aggregate count of patients matching the query instead. (bottom). HLL + Shuffle. We do a coordinated random shuffling so the central server does not know what the original bucket leading 1-indicators were.

## III. DISCUSSION

Here, we have surveyed and benchmarked a range of methods, exploring the trade-offs in privacy, accuracy, and speed. We explicitly do not endorse a single one-size-fits-all method, as different applications and institutions will have different needs. Indeed, we envision that in practice, queries will be first run using a fast, private method (such as the currently standard aggregate counts, perhaps combined with MPC); given those rough results and the needs of the researcher, one of the more accurate HLL-based methods can then be used if more accuracy is desired. In the final stage of research (e.g. in preparation for a full clinical trial), institutions can then sign the necessary data use agreements to share raw identifiable information. We believe that as federated data networks expand to include more institutions and data types (clinical, genomic, environmental, etc.), researchers will increasingly depend on fast, accurate and secure query tools to get the greatest possible scientific value from the networks.

## IV. METHODS

We consider several variants of the base methods (hashed IDs, aggregate count, secure sharing, and HLL) that further protect patient privacy. Below, we first describe each algorithm for querying the network. Then, we define a privacy risk score and provide details on how we benchmark the algorithms using simulated hospital networks.

### A. Query Method + obfuscation details

Here, we describe in detail the protocols we compared, as well as several that we did not directly compare against for reasons that will be explained. The basic model will assume that the researcher sending the query goes through one of the participating hospitals in the network, and asks a query of the form: “how many patients have condition X across the hospital network?”

#### 1) Count

The query is sent from a hospital to the hub. The hub broadcasts out that query to each hospital. Each hospital runs the query locally, producing a count, which is then sent back to the hub. The hub returns two numbers to the originating hospital: (1) the maximum count from a hospital and (2) the sum of counts from all hospitals. The former corresponds to a lower bound on the result, because even in the event of significant overlapping patients between hospitals, there are at least as many unique patients across the network as there are at a single hospital. The latter summation is obviously an upper bound, though it might be a substantial overestimate when there is significant overlap between hospitals.

#### 2) Count + Mask

The procedure is identical to Count, except that if the actual count of a hospital is between 1 through 9 inclusive, the hospital returns 10 to the hub instead. This masking procedure ensures that no non-zero number corresponds to fewer than 10 patients, ensuring 10-anonymity. Both the PCORNet and ACT networks use Count + Mask. ACT further obfuscates the result by adding a small random number between −10 and +10 to the actual count [Murphy 2002]; though, we ignore this in our analyses.

#### 3) Count + MPC

This protocol is based on the ElGamal cryptosystem [ElGamal 1985] using a distributed private key to ensure that no one party can decrypt intermediate data, secure in the Honest But Curious even if all hospitals but one and the hub are compromised. The major disadvantage is that the MPC requires all hospitals to respond before any answer can be given. In large networks, it is likely that some hospitals will either be slow to respond or not respond at all [Weber 2015], which limits this protocol to only small networks in practice.

We take advantage of the multiplicatively homomorphic property of ElGamal encryption, which means that given an encryption function *E* and decryption function *D, D*(*E*(*ab*)) = *ab*. It is secure in the semi-honest framework assuming the difficult of the discrete logarithm problem. The algorithm has three parts (key generation, encryption, and decryption), which requires two rounds of communication between the hospitals and the hub.

##### Key generation

We use a fixed 1024-bit prime *p* and appropriate generator *g* as the basis of all our cryptographic keys (see Supplementary Information: ElGamal constants). All arithmetic will be performed in that prime field defined by *p* (i.e. will be performed modulus *p*).

Given *p* and *g*, a private/public keypair consists of a random number *x* ∈ [2, *p* − 1] as the private key, and *y* = *g*^*x*^. We wish to ensure that no one party ever has access to *x*, so we use a distributed key generation protocol. Each hospital *i* generates a random *x*_*i*_ and produces the corresponding 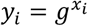, which it sends to the central hub. The hub then computes *y* = ∏*y*_*i*_. Note that 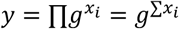, so *y* is the public key corresponding to the private key *x* = ∑*x*_*i*_, but no hospital actually knows the summed value *x*. The hub returns the public key *y* to each hospital.

##### Encryption

The standard ElGamal encryption function *E*(*m*) = (*g*^*z*^, *my*^*z*^), where *z* in a random integer in the field. Note that ElGamal has the nice property that the same plaintext message will be encrypted to a many possible encrypted ciphertexts because of *z*. This is essential to defeating dictionary attacks on the ciphertext. As a technical note, in order to have provable security, the message *m* must be a quadratic residue of the field, which we will ensure the in the protocol described.

##### Decryption

The standard ElGamal decryption function 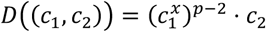 Note that we can do a distributed decryption for a ciphertext by sending each hospital *c*_1_, and asking the hospitals to return 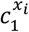, which when multiplied together give 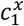.

##### Round 1: encryption and summation

Each hospital runs the query locally, producing a count *a*_*i*_, and then sends the value 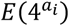 back to the hub (we use 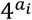 because it is a quadratic residue). The hub computes 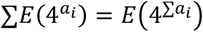.

##### Round 2: decryption

The encrypted sum is decrypted using the distributed decryption protocol described above, giving 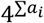. Of course, performing discrete logarithms is hard (or else ElGamal encryption would not be secure), but we can precompute a discrete log table for powers of 4 relatively easily. Using that lookup table, the hub produces ∑*a*_*i*_ as the final response for the query.

#### 4) HyperLogLog

Upon receiving the query, a hospital generates a list of patient IDs that match the query. Each hospital needs to use the same ID for the same patient. Because there is no universal patient identifier (UPI), the ID should be based on information likely to be unique to the patient and available at all hospitals, such as the concatenation of the patient’s first name, last name, and date of birth [Grannis 2002]. (See “Supplementary Information: Generating Patient IDs” for additional details and limitations of generating a patient ID.)

The hospital runs all of the patient IDs through SHA-1, producing a 160-bit pseudorandom number. The first 64-bits are interpreted as an integer *B*, and the patient is put into bucket *B* % *k*, where *k* is the number of buckets. The hospital then finds the position *V* of the first bit set to 1 in bits 65-128 of the SHA1-string. Within each bucket, the hospital stores the largest value *V* corresponding to a patient. The list of bucket values is the HLL sketch from that hospital.

The hospitals send those HLL sketches to the central hub. The hub combines the sketches by taking the maximum within each bucket across hospital sketches, generating the sketch of the union. The hub then estimates the cardinality *C* of the union sketch using the standard HLL estimator [Flajolet, 2007]. The hub also provides a 95% confidence interval by using the fact that the standard deviation of the estimate is around 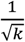, so 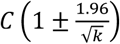 gives the lower and upper bounds of a 95% confidence interval.

Note that although we described the full procedure above as a linear process starting with the receipt of the query, the hospital can precompute the buckets *B* and values *V* for all of their patients, reducing the time needed to generate the HLL sketch for given query.

#### 5) HyperLogLog + Mask

This procedure is identical to HLL, except that the hospital precomputes a list of bucket values that are less than 10-anonymous. If after generating the HLL sketch corresponding to the query, a hospital sees that there is a bucket that is not 10-anonymous, the hospital aborts and reverts to the count + Mask-10 algorithm, where only a single (possibly masked) aggregate count is returned.

The hub thus receives a combination of sketches and masked counts. The hub combines together the sketches using the HLL cardinality estimator to get an estimate of the count of the union of all the hospitals that sent sketches with appropriate 95% error bounds. From that, the hub goes through something similar to the Count-procedure. The hub returns two numbers: the sum of all raw hospital counts plus the 95% confidence interval maximum for the HLL union count, which gives an upper bound, and the maximum of the set of raw counts or the 95% confidence interval minimum for the HLL union, which gives a lower bound.

#### 6) HyperLogLog + Shuffle

When a hospital sends a query to the hub, it sends both a query, and a random string encrypted with public keys of each of the other hospitals in the network (using any kind of standard off-the-shelf asymmetric key encryption, as used in protocols like RSA, HTTPS, etc.). The hub forwards the query to all the hospitals, plus the encrypted random string, which the hub itself cannot decrypt. Each hospital then uses an ordinary HLL sketch, but shuffles the ordering of the buckets using the random string to determine the sort order, and then sends the shuffled sketch to the hub.

Because every hospital performs the same permutation, the sketches can still be combined and the normal estimators used. However, the hub, without knowing the random string, cannot know which bucket was which. Normally, an HLL bucket is less than 10-anonymous if that value + bucket pair corresponds to fewer than 10 individuals at the hospital. With shuffling, an HLL bucket is less than 10-anonymous only if that value corresponds to fewer than 10 individuals at the hospital. On average, this decreases the risk by dividing the risk score by the number of buckets. In other words, the buckets partition the patient population into smaller, more identifiable groups. By shuffling the buckets, it is no longer known which partition the value came from, which makes the value less identifiable.

#### 7) HyperLogLog + Rehash

This procedure is similar to that of HyperLogLog + Shuffle. However, instead of using the random string to shuffle buckets, the hospitals completely regenerate the HLL sketch while prepending that random string to the patient IDs before hashing using SHA-1. This procedure takes more time, but also means that the hub cannot use a dictionary attack at all, because it does not know the random string. Thus, all patients are guaranteed 10-anonymity if the random string is not revealed to the hub.

#### 8) HyperLogLog + MPC

Like Count + MPC, this method is based off of the ElGamal homomorphic cryptosystem, and we use the same primitives as in that method (with the same security guarantees). We additionally take inspiration from a previous paper applying MPC to a Flajolet-Martin style approximate counter [Dong 2017]. The key setup and exchange are identical to Count + MPC, as well as the encryption and decryption routines, so we only describe the following rounds:

##### Round 1: encryption and merging

Each hospital begins by generating an HLL sketch of the query. We then unroll each bucket *B*_*j*_ = *v*_*j*_ of the sketch into a binary string of length 32 with *v*_*j*_1’s and 32 − *v*_*j*_ 0’s. i.e. if *v*_*j*_ = 10, the binary string would be “11111111110000000000000000000000”. However, ElGamal homomorphic encryption is only secure when using non-zero quadratic residues of the prime field. So we turn that string into a vector, replacing 1’s with 4’s and 0’s with 1’s, resulting in a vector of length 32, [4, 4, 4, 4, 4, 4, 4, 4, 4, 4, 1, …, 1]. The hospital then encrypts each of these unrolled bucket vectors into [*E*(4), …, *E*(4), *E*(1), … *E*(1)], and send them to the hub. Note that we rely the fact that ElGamal encryption is probabilistic, so each of the 4’s encrypts to a different ciphertext, and so do each of the 1’s. Thus, the encrypted vector does not reveal any information about the underlying binary bitstring.

The hub receives the encrypted HLL sketches from each hospital, and then takes the product across hospitals of each position in the unrolled bucket vectors, giving a product vector [∏*x*_1,*i*_, …, ∏*x*_32,*i*_]. Because ElGamal is multiplicatively homomorphic, ∏*x*_1,*i*_ = *E*(1) if and only if all *x*_*j,i*_ = *E*(1). Were we to decrypt this vector, it would reveal the maximum bucket value for this bucket, because the vector would be equal to 1 at all indices above that value. However, this leaks information because the other indices would have some value 4^*y*^, where *y* is the number of times a hospital had a value of at least that index.

To resolve this information leakage, we use a private equality test [Jakobsson & Juels, 2000]. Given two ciphertexts (*c*_1_, *c*_2_) and 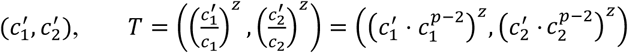, where *z* is a random integer, is a private equality test. More precisely, *D*(*T*) = 1 if and only if 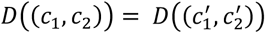. More importantly, *D*(*T*) is a random integer (different from *z*) if the two ciphertexts were not equal in the plaintext space. The hub thus does a private equality test of all the combined encrypted bucket values, testing if they are equal to 1, and masking the result if they are not equal to 1. Those new masked vectors do not leak any information, revealing only the maximum value of the bucket across hospitals.

##### Round 2: decryption

We now run the distributed decryption protocol on each of those masked vector elements. Because each element is independent, they can be decrypted in parallel in only one round of communication. For each bucket, the hub then looks at the maximum index that is not equal to 1, which corresponds to the maximum bucket value across hospitals; this procedure allows the hub to reconstruct the merged HLL sketch. Once given a merged HLL sketch, the hub can then follow the rest of the standard procedure for the HyperLogLog method.

#### 9) HyperLogLog + Shuffle + MPC

This procedure is simply a combination of HyperLogLog + Shuffle and HyperLogLog + MPC. Each hospital simply shuffles their buckets according to the random string prior to performing encryption. The rest of the procedure is identical to HyperLogLog + MPC.

#### 10) HashedIDs

The query is sent from a hospital to the hub. The hub broadcasts out that query to each hospital. Each hospital runs the query locally, producing a list of matching patient IDs. (Patient IDs are generated the same way as in HyperLogLog. See Supplementary Information.). The patient IDs are then hashed via SHA-1. That list of hashed IDs is then sent back to the hub. The hub then deduplicates the list (e.g. via a hash table), and counts the number of unique individuals matching the query across the entire hospital system. That count is returned as the (exact) answer to the query.

Note that the list of hashed IDs can be precomputed, just as in HyperLogLog, because each patient’s hashed ID will not change. This also means of course that a dictionary attack by the hub has a high likelihood of success.

#### 11) HashedIDs + Rehash

This is identical to HashedIDs, except that the originating hospital also sends a random string encrypted with the public keys of each of the other hospitals. Each hospital rehashes all the patients, prepending the random string before running it through SHA-1. By doing so, because the hub does not know the random prefix string, it cannot do a dictionary attack to reverse the hash function, and thus all patients get 10-anonymity. Of course, rehashing all patients takes additional computational time.

#### 12) Secure methods that are not scalable to large networks

Above, we described protocols that we quantitatively benchmark in this study, including two secure MPC protocols we implemented. Count + MPC is just a straight-forward implementation of secure MPC summation, but HLL + MPC is a protocol we developed ourselves, inspired by Dong, et al [2017]. The reason we developed that protocol instead of using existing protocols from the cryptographic literature is that most such methods are impractically slow, due to bad scaling of communication and computation requirements. Here, we describe a few secure MPC protocols that provide privacy guarantees without the need for a trusted 3^rd^ party. However, because secure MPC and homomorphic encryption are computationally complex, they could take on the order of days to weeks for a single query in a large network. This makes them impractical except for very small networks. As a result, we do not include them in our benchmarking simulations.

One MPC approach is to use a pairwise private intersection protocol [Kolesnikov 2017, de Christofaro 2012], which securely determines the number of shared patients between two sites. Subtracting this from the sum of the counts from each site gives the total number of distinct patients. However, the number of required pairwise and multi-way comparisons grows exponentially with the number of sites, making this impractical for large networks. Patient partitioning [Weber 2013] and cryptosets [Swamidass 2015] are related non-MPC methods that have similar scalability problems due to the number of patient slices. A recent approach using counting Bloom filters is able to solve the deduplication problem without pairwise comparisons, but due to the nature of Bloom filters scales linearly in the number of patients and requires at least two trusted data custodians even in a semi-honest framework [Yigzaw 2017].

Other work in the MPC literature has produced algorithms for directly computing unions and deduplications of sets without the problem of exponential comparisons. Unfortunately, this comes at the cost of either significantly more computation time and communication bandwidth requirements, which can be on the order of gigabytes of shared data for a single query, with linear communication complexity and super-linear time-complexity in the number of patients [Fenske 2017]. A more recent approach combines a Flajolet-Martin style estimator with a secure MPC protocol [Dong 2017]. The algorithm has logarithmic space complexity, in that the number of bits needed scales logarithmically with respect to the number of patients who match the query. However, the trade-off is that it requires numerous back-and-forth communication---on the order of log *N* rounds, where *N* is the total patient population---between all the hospitals in the network to execute the protocol. As mentioned above though, our HLL + MPC protocol is heavily based off of Dong, et al.

The root of the issue is that in the context of a federated network of hospitals, if each hospital acts as a computing party for an MPC protocol, then each hospital can guarantee to itself that at least it itself is not malicious. This feature is desirable for hospitals, because it means they do not have to trust anyone but themselves. However, most MPC methods scale badly in the number of computing parties; using semi-trusted dedicated compute parties can help, but that still requires trusting those compute parties to not collude. In recent years, more scalable secure MPC protocols have been introduced to solving distributed genome-wide association studies [Cho, 2018] and pharmacological collaborations [Hie & Cho, 2018], but these protocols are not practical for the near-real-time results that clinical researchers expect (indeed, in that context, it is considered fast to get results in weeks). For this reason, we only compared against the two MPC protocols we ourselves implemented, which are designed to be scalable at the level we need for clinical queries.

### B. Privacy risk score – m-anonymity

We define a piece of aggregate information, or statistic, as less than *m*-anonymous if it includes at least 1 individual and could have been generated by fewer than *m* individuals in some background population. As long as patients have 2-anonymity, they have not been fully revealed. However, in practice, hospitals are usually more conservative. One study recommended 5-anonymity for hospitals [Emam 2008], but the national PCORNet and ACT networks go even higher, requiring 10-anonymity. For purposes of this paper, we will use 10-anonymity throughout our analysis to be consistent with these existing networks. We will define the privacy risk for a release of data as the number of statistics revealed to the adversary that are not 10-anonymous.

For the background population we use the patient population at a hospital, because the hub generally knows when a piece of information comes from a particular hospital. In the case where we use MPC to merge data across the network, however, the background population can be taken to be the patient population across the entire network, as no one party sees the information from a single hospital.

In the case of a single count from a hospital, whether or not that count is 10-anonymous is easy to determine: if the count is between 1 and 9 inclusive, then it is not 10-anonymous; else, it is. Note that this is not a perfect proxy, because while a single count may be 10-anonymous, multiple counts from the same hospital might not be. For example, if the count of male patients is 10 and the count of male + female patients is 11, then two counts, while individually 10-anonymous, can together be combined to reveal that there is only 1 female patient. Although here we analyze only the privacy risk from revealing a single count from a hospital, so we do not worry about that, it is still worth remembering that even aggregate counts >10 are not perfect 10-anonymity.

For a hashed value generated from a patient ID, we consider it 10-anonymous if the adversary cannot reverse the hash function to figure out the original patient ID to within 10 patients. Luckily, cryptographic hash functions are one-way, meaning that the function cannot be directly reversed. Unfortunately, since the space of patient IDs (e.g. social security numbers) is constrained, an adversary can simply create a rainbow table of the hashed values of every possible patient ID, and then simply do a lookup. Thus, a hashed value is only 10-anonymous if at least 10 patients in the background population hash to that particular value. Unfortunately, for hashed IDs that are sufficiently large to do deduplication of patients (e.g. 32- or 64-bits), the very property that allows deduplication also ensures that close to none of the hashed IDs are 10-anonymous.

HyperLogLog buckets can be thought of as a much shorter hashed ID. Whereas we might use a 64-bit hash when using Hashed IDs, the HyperLogLog bucket stores only the position of the first 1 bit in that 64-bit hash. This increases the number of collisions considerably. An HLL bucket with value *x* is 10-anonymous if at least 10 patients in the background population have hashes where the leading 1-indicator indicator is in position *x*, which happens much more often. Additionally, there are generally many fewer HLL buckets than patient IDs, so fewer potentially risky statistics are revealed to begin with.

As an aside, our privacy risk analysis differs considerably from Desfontaines, et al (2018), who argue that “cardinality estimators do not preserve privacy.” However, their threat model includes an adversary who has incremental access to the sketches as they are being generated, rather than only a single sketch per hospital for a query. Were a hospital compromised to the point where their internal systems were constantly revealing incremental sketch updates, the privacy loss from HyperLogLog would be the least of their worries.

### C. Benchmarking

#### 1) Simulating a geographic hospital network

Because of patient privacy, we cannot test the algorithms using actual hospital data. We therefore built a simulation of a set of hospitals spread geographically with highly varying sizes and overlap.

First we model geographic spread by placing 100 cities uniformly randomly in a 2D unit square. City sizes are often modeled to have lognormal distributions [Berry 1961], with probability density function

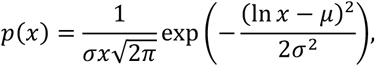

where *μ* and *σ* are respectively the mean and standard deviation of the underlying normal distribution.

Each of the 100 hospitals in our network is assumed to draw primarily from one of those cities. We randomly sample 100 numbers from a lognormal distribution with *μ* = 0 and *σ* = 1.2, and then scale up all the numbers such that the sum is 100 million total unique patients. Each patient is assigned a number between 1 and 100 million, and is then placed in one of the 100 hospitals as their home hospital, according to the scaled up lognormal size distribution computed earlier.

For each patient, we draw a random integer from a binomial distribution 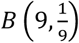, which will denote the number of additional hospitals patients are assigned to. Here the intuition is that most patients are only at a single hospital, but some patients are admitted to many hospitals. However, by choosing those parameters of the binomial distribution, we ensure that on average, patients are admitted to 2 hospitals (their home hospital, and one additional one as the mean of the binomial distribution is 1).

Then we assume that patients who are admitted to multiple hospitals are more likely to go to nearby ones, according to the hospital locations in the unit square we assigned earlier. We assume that the probability that a patient chooses a particular additional hospital is inversely proportional to the square of the distance between the new hospital and the patient’s home hospital. Using this probability distribution, we assign each patient to their additional hospitals.

By using this procedure, we generate hospitals that start with lognormal sizes, following city size distributions, but with some smoothing of the sizes because some patients will go to multiple hospitals.

#### 2) Benchmarking methodology

All benchmarks were run using Python code available at https://github.com/yunwilliamyu/secure-distributed-union-cardinality. The benchmarks were run on an 8-core AMD Ryzen 1700 CPU with 16 GiB of RAM running Ubuntu 18.04.2 LTS. We measured wall-clock time for each pipeline component for time-complexity, and serialized bytestrings in each communication round for transmission space-complexity. Methods analyzed were aggregate counts, HyperLogLog (HLL) sketches, and hashed IDs, paired with various obfuscation techniques of masking, rehashing, shuffling, and MPC. Note that we explore different values for the number of buckets, and title the method by the number of bits used for that bucket (so HLL-7 means we use 2^7^ = 128 buckets). We simulate hospital networks with 100 million total unique patients, distributed across 100 geographically separated hospitals, with each patient on average appearing at 2 hospitals (though individual patients might appear at more or fewer hospitals), and being more likely to appear at nearby hospitals (as specified in the previous section).

In Supplementary Table 1, we give the computational and communication costs. We give the empirical scalings of runtime and transmission bandwidth from the experiments we run. By combining the two, we can provide an upper bound on the added CPU and transmission costs from using the various query methods and obfuscations.

We then run queries matching 1, 10, 100, 1 thousand, 10 thousand, 100 thousand, 1 million, 10 million, or 100 million patients using the different methods. Mean wait time is the average hospital computation time + hub computation time. Max wait time is the maximum hospital computation time for a run + hub computation time. To measure accuracy, we provide 95 percent confidence intervals based off 100 simulated experiments and compared them against the true number of patients matching the query. To measure privacy, we count the number of statistics (i.e. a count, HLL bucket, or hash) without 10-anonymity revealed to either the hub, or the hub colluding with a hospital (Supplementary Tables 2-10).

## V. ACKNOWLEDGMENT

This study was supported by National Institutes of Health (NIH) Big Data to Knowledge (BD2K) awards U54HG007963 from the National Human Genome Research Institute (NHGRI) and U01CA198934 from the National Cancer Institute (NCI). Y.W.Y. was also supported by training grant T15LM007092 from the NIH NLM.

## VI. AUTHOR CONTRIBUTIONS

G.M.W. and Y.W.Y. conceived the project and wrote the manuscript. Y.W.Y. designed and implemented the software, and also performed the mathematical analyses. G.M.W. guided the direction of the research and provided critical advice on the constraints of the task.

