## Supplemental Information for "Federated queries of clinical data repositories: balancing accuracy and privacy"

### Generating patient IDs

For both the HyperLogLog and HashedIDs methods, a unique identifier (ID) is needed for each patient. Each hospital needs to use the same ID for the same patient. Because there is no universal patient identifier (UPI), the ID should be based on information likely to be unique to the patient and available at all hospitals, such as the concatenation of the patient's first name, last name, and date of birth. However, there are limitations to this because of missing data, data errors (e.g., a misspelled name), and data that change over time (e.g., a person changes his or her name).

The best combination of demographic variables depends on whether it is preferable for queries to over- or under-count the number of matching patients [Grannis 2002]. For example, using just first initial plus last name will cause many people to have the same ID; while, combining first and last name, date of birth, social security number, and zip code increases the chance that the same person has different IDs at different hospitals.

Although it would increase query time and privacy risk, a hospital network can generate multiple IDs for each patient, using different combinations of variables. The same query can be run once for each ID, producing a range of results. The researcher can use this to gauge what the true number of matching patients might be.

### ElGamal constants

The 1024-bit prime used is:

```
980280362726590313715607422429551698619105506348111713
034013583788161326369921156306749439948292232187834981
114813830512407606566691503608138648619957629317511318
723406340506422747334388220492377414235316985427964841
483121858791867814907317712291782727600552816246987650
8421706801816828847677451793907439076 7
```

The generator we use is:

```
242064085869641024218127655172409731313195116710442028
562029920756038984848088822296764302596885435248284967
249404984657383450832959107783541977139869194907266271
150046901293736765904796968178129569636323302524909947
269844370371135196758871673117158639233223412811927551
80251058938103973809860137264177781321
```

### Benchmark results summary

| Algorithm | | Added Time Complexity | | | Actual Additional Time (s) | | Actual Time Examples (s) | | Transfer Time | | Actual Data |
|---|---|---|---|---|---|---|---|---|---|---|---|
| Method | Obfuscation | Sketch | Obfuscate | Hub | Site | Hub | 1 Patient | 100M Patients | Rounds | Bytes | 100M Patients |
| Count | | | | O(n) | | $10^{-7}n$ | 0.000 | 0.000 | 1 | $4n$ | 400 bytes |
| Count | Mask | | O(1) | O(n) | 10^-7 | $10^{-7}n$ | 0.000 | 0.000 | 1 | $4n$ | 400 bytes |
| Count | MPC | | O(1) | O(n) | 0.049 | $0.05 + 0.006n$ | 0.100 | 0.100 | 3 | $896n$ | 87.5 KiB |
| HLL | | O(k) | | O(nk) | | $5 \times 10^{-9} \cdot nk$ | 0.006 | 0.005 | 1 | $nk$ | 12800 bytes |
| HLL | Mask | O(k) | O(k) | O(nk) | $10^{-7} \cdot k$ | $5 \times 10^{-9} \cdot nk$ | 0.007 | 0.001 | 1 | $nk$ | 12800 bytes |
| HLL | Shuffle | O(k) | O(k) | O(nk) | $10^{-7} \cdot k$ | $5 \times 10^{-9} \cdot nk$ | 0.006 | 0.005 | 1 | $nk$ | 12800 bytes |
| HLL | Rehash | O(k) | O(m) | O(nk) | $6.5 \times 10^{-6} \cdot m$ | $5 \times 10^{-9} \cdot nk$ | 0.007 | 8.757 | 1 | $nk$ | 12800 bytes |
| HLL | MPC | O(k) | O(k) | O(nk) | $0.05k$ | $0.00235nk + 0.197k + 0.027$ | 40.930 | 41.010 | 3 | $256n + 20480nk$ | 250 MiB |
| HLL | Shuffle+MPC | O(k) | O(k) | O(nk) | $0.05k$ | $0.00235nk + 0.197k + 0.027$ | 40.930 | 41.010 | 3 | $256n + 20480nk$ | 250 MiB |
| HashedIDs | | | | O(mn) | | $10^{-7} \cdot mn$ | 0.003 | 12.670 | 1 | $4m$ | 8 MB |
| HashedIDs | Rehash | | O(m) | O(mn) | $6.5 \times 10^{-6} \cdot m$ | $10^{-7} \cdot mn$ | 0.003 | 14.350 | 1 | $4m$ | 8 MB |

*$n$=number of hospitals, $m$=patients per hospital, $k$=number of HLL buckets

**Supplementary Table 1**. Time and Space complexity for various methods. These are upper bounds on the added compute time when using a particular method + obfuscation. We also have the space-complexity of the amount of data that the hospitals and hub have to send over the network. Note that actual times given are based on assuming that the hospitals run their local computations in parallel.

### Benchmark results details

| Method | Obfuscation | Estimated Count | | Wait (s) | | Risk - Hub | Risk - Hub + Hospital |
|---|---|---|---|---|---|---|---|
| | | Absolute | Relative | Mean | Max | | |
| Count | None | [1.000, 3.525] | [0%, 252%] | 0 | 0 | 1.91 | 1.91 |
| | Mask | [10.000, 35.25] | [900%, 3425%] | 0 | 0 | 0 | 0 |
| | MPC | [1.000, 3.525] | [0%, 252%] | 0.099 | 0.099 | 0 | 0 |
| HLL0 | None | [0.694, 22.20] | [-31%, 2120%] | 0 | 0 | 0 | 0 |
| | Shuffle | [0.694, 22.20] | [-31%, 2120%] | 0 | 0 | 0 | 0 |
| | Rehash | [0.694, 22.20] | [-31%, 2120%] | 0 | 0 | 0 | 0 |
| | Mask | [0.694, 22.20] | [-31%, 2120%] | 0 | 0 | 0 | 0 |
| | MPC | [0.694, 22.20] | [-31%, 2120%] | 0.385 | 0.385 | 0 | 0 |
| | Shuffle+MPC | [0.694, 22.20] | [-31%, 2120%] | 0.385 | 0.385 | 0 | 0 |
| HLL1 | None | [1.386, 1.386] | [39%, 39%] | 0 | 0 | 0 | 0 |
| | Shuffle | [1.386, 1.386] | [39%, 39%] | 0 | 0 | 0 | 0 |
| | Rehash | [1.386, 1.386] | [39%, 39%] | 0 | 0 | 0 | 0 |
| | Mask | [1.386, 1.386] | [39%, 39%] | 0 | 0 | 0 | 0 |
| | MPC | [1.386, 1.386] | [39%, 39%] | 0.675 | 0.675 | 0 | 0 |
| | Shuffle+MPC | [1.386, 1.386] | [39%, 39%] | 0.675 | 0.675 | 0 | 0 |
| HLL4 | None | [1.033, 1.033] | [3%, 3%] | 0.001 | 0.001 | 0 | 0 |
| | Shuffle | [1.033, 1.033] | [3%, 3%] | 0.001 | 0.001 | 0 | 0 |
| | Rehash | [1.033, 1.033] | [3%, 3%] | 0.001 | 0.001 | 0 | 0 |
| | Mask | [1.033, 1.033] | [3%, 3%] | 0.001 | 0.001 | 0 | 0 |
| | MPC | [1.033, 1.033] | [3%, 3%] | 4.781 | 4.781 | 0 | 0 |
| | Shuffle+MPC | [1.033, 1.033] | [3%, 3%] | 4.781 | 4.781 | 0 | 0 |
| HLL7 | None | [1.004, 1.004] | [0%, 0%] | 0.006 | 0.006 | 0 | 0 |
| | Shuffle | [1.004, 1.004] | [0%, 0%] | 0.006 | 0.006 | 0 | 0 |
| | Rehash | [1.004, 1.004] | [0%, 0%] | 0.007 | 0.007 | 0 | 0 |
| | Mask | [1.004, 1.004] | [0%, 0%] | 0.007 | 0.007 | 0 | 0 |
| | MPC | [1.004, 1.004] | [0%, 0%] | 37.73 | 37.73 | 0 | 0 |
| | Shuffle+MPC | [1.004, 1.004] | [0%, 0%] | 37.73 | 37.73 | 0 | 0 |
| HLL15 | None | [1.000, 1.000] | [0%, 0%] | 1.463 | 1.463 | 0.31 | 0.31 |
| | Shuffle | [1.000, 1.000] | [0%, 0%] | 1.463 | 1.463 | 0 | 0.31 |
| | Rehash | [1.000, 1.000] | [0%, 0%] | 1.623 | 1.647 | 0 | 0.31 |
| | Mask | [1.000, 21.00] | [0%, 2000%] | 1.632 | 1.632 | 0 | 0 |
| Hashed IDs | None | [1.000, 1.000] | [0%, 0%] | 0.001 | 0.001 | 1.91 | 1.91 |
| | Rehash | [1.000, 1.000] | [0%, 0%] | 0.001 | 0.001 | 0 | 1.91 |

**Supplementary Table 2**. 1 patient query. We measure the accuracy, wait time, and privacy risk of running the query using raw counts, HyperLogLog sketches of various sizes, and sending full Hashed patient IDs.

| Method | Obfuscation | Estimated Count | | Wait (s) | | Risk - Hub | Risk - Hub + Hospital |
|---|---|---|---|---|---|---|---|
| | | Absolute | Relative | Mean | Max | | |
| Count | None | [1.475, 24.00] | [-85%, 140%] | 0 | 0 | 15.72 | 15.72 |
| | Mask | [10.000, 205.2] | [0%, 1952%] | 0 | 0 | 0 | 0 |
| | MPC | [15.47, 24.00] | [55%, 140%] | 0.099 | 0.099 | 0 | 0 |
| HLL0 | None | [2.776, 88.82] | [-72%, 788%] | 0 | 0 | 0 | 0 |
| | Shuffle | [2.776, 88.82] | [-72%, 788%] | 0 | 0 | 0 | 0 |
| | Rehash | [2.776, 88.82] | [-72%, 788%] | 0 | 0 | 0 | 0 |
| | Mask | [2.776, 88.82] | [-72%, 788%] | 0 | 0 | 0 | 0 |
| | MPC | [2.776, 88.82] | [-72%, 788%] | 0.385 | 0.385 | 0 | 0 |
| | Shuffle+MPC | [2.776, 88.82] | [-72%, 788%] | 0.385 | 0.385 | 0 | 0 |
| HLL1 | None | [3.332, 35.23] | [-67%, 252%] | 0 | 0 | 0 | 0 |
| | Shuffle | [3.332, 35.23] | [-67%, 252%] | 0 | 0 | 0 | 0 |
| | Rehash | [3.332, 35.23] | [-67%, 252%] | 0 | 0 | 0 | 0 |
| | Mask | [3.332, 35.23] | [-67%, 252%] | 0 | 0 | 0 | 0 |
| | MPC | [3.332, 35.23] | [-67%, 252%] | 0.675 | 0.675 | 0 | 0 |
| | Shuffle+MPC | [3.332, 35.23] | [-67%, 252%] | 0.675 | 0.675 | 0 | 0 |
| HLL4 | None | [5.995, 15.69] | [-40%, 57%] | 0.001 | 0.001 | 0 | 0 |
| | Shuffle | [5.995, 15.69] | [-40%, 57%] | 0.001 | 0.001 | 0 | 0 |
| | Rehash | [5.995, 15.69] | [-40%, 57%] | 0.001 | 0.001 | 0 | 0 |
| | Mask | [5.995, 15.69] | [-40%, 57%] | 0.001 | 0.001 | 0 | 0 |
| | MPC | [5.995, 15.69] | [-40%, 57%] | 4.778 | 4.778 | 0 | 0 |
| | Shuffle+MPC | [5.995, 15.69] | [-40%, 57%] | 4.778 | 4.778 | 0 | 0 |
| HLL7 | None | [8.261, 10.41] | [-17%, 4%] | 0.006 | 0.006 | 0.01 | 0.01 |
| | Shuffle | [8.261, 10.41] | [-17%, 4%] | 0.006 | 0.006 | 0 | 0.01 |
| | Rehash | [8.261, 10.41] | [-17%, 4%] | 0.007 | 0.007 | 0 | 0 |
| | Mask | [8.261, 10.41] | [-17%, 4%] | 0.007 | 0.007 | 0 | 0 |
| | MPC | [8.261, 10.41] | [-17%, 4%] | 37.72 | 37.72 | 0 | 0 |
| | Shuffle+MPC | [8.261, 10.41] | [-17%, 4%] | 37.72 | 37.72 | 0 | 0 |
| HLL15 | None | [10.00, 10.00] | [0%, 0%] | 1.477 | 1.477 | 4 | 4 |
| | Shuffle | [10.00, 10.00] | [0%, 0%] | 1.477 | 1.477 | 0 | 4 |
| | Rehash | [10.00, 10.00] | [0%, 0%] | 1.636 | 1.659 | 0 | 3.8 |
| | Mask | [10.000, 87.53] | [0%, 775%] | 1.578 | 1.578 | 0 | 0 |
| Hashed IDs | None | [10.000, 10.000] | [0%, 0%] | 0.002 | 0.002 | 19 | 19 |
| | Rehash | [10.000, 10.000] | [0%, 0%] | 0.002 | 0.002 | 0 | 19 |

**Supplementary Table 3**. 10 patient query. We measure the accuracy, wait time, and privacy risk of running the query using raw counts, HyperLogLog sketches of various sizes, and sending full Hashed patient IDs.

| Method | Obfuscation | Estimated Count | | Wait (s) | | Risk - Hub | Risk - Hub + Hospital |
|---|---|---|---|---|---|---|---|
| | | Absolute | Relative | Mean | Max | | |
| Count | None | [9.000, 205.6] | [-91%, 106%] | 0 | 0 | 59.78 | 59.78 |
| | Mask | [10.000, 734.1] | [-90%, 634%] | 0 | 0 | 0 | 0 |
| | MPC | [176.9, 205.6] | [77%, 106%] | 0.099 | 0.099 | 0 | 0 |
| HLL0 | None | [11.10, 541.8] | [-89%, 442%] | 0 | 0 | 0 | 0 |
| | Shuffle | [11.10, 541.8] | [-89%, 442%] | 0 | 0 | 0 | 0 |
| | Rehash | [11.10, 541.8] | [-89%, 442%] | 0 | 0 | 0 | 0 |
| | Mask | [11.10, 541.8] | [-89%, 442%] | 0 | 0 | 0 | 0 |
| | MPC | [11.10, 541.8] | [-89%, 442%] | 0.384 | 0.384 | 0 | 0 |
| | Shuffle+MPC | [11.10, 541.8] | [-89%, 442%] | 0.384 | 0.384 | 0 | 0 |
| HLL1 | None | [19.99, 807.6] | [-80%, 708%] | 0 | 0 | 0 | 0 |
| | Shuffle | [19.99, 807.6] | [-80%, 708%] | 0 | 0 | 0 | 0 |
| | Rehash | [19.99, 807.6] | [-80%, 708%] | 0 | 0 | 0 | 0 |
| | Mask | [19.99, 807.6] | [-80%, 708%] | 0 | 0 | 0 | 0 |
| | MPC | [19.99, 807.6] | [-80%, 708%] | 0.676 | 0.676 | 0 | 0 |
| | Shuffle+MPC | [19.99, 807.6] | [-80%, 708%] | 0.676 | 0.676 | 0 | 0 |
| HLL4 | None | [58.08, 158.6] | [-42%, 59%] | 0.001 | 0.001 | 0 | 0 |
| | Shuffle | [58.08, 158.6] | [-42%, 59%] | 0.001 | 0.001 | 0 | 0 |
| | Rehash | [58.08, 158.6] | [-42%, 59%] | 0.001 | 0.001 | 0 | 0 |
| | Mask | [58.08, 158.6] | [-42%, 59%] | 0.001 | 0.001 | 0 | 0 |
| | MPC | [58.08, 158.6] | [-42%, 59%] | 4.782 | 4.782 | 0 | 0 |
| | Shuffle+MPC | [58.08, 158.6] | [-42%, 59%] | 4.782 | 4.782 | 0 | 0 |
| HLL7 | None | [87.68, 115.3] | [-12%, 15%] | 0.006 | 0.006 | 0.07 | 0.07 |
| | Shuffle | [87.68, 115.3] | [-12%, 15%] | 0.006 | 0.006 | 0 | 0.07 |
| | Rehash | [87.68, 115.3] | [-12%, 15%] | 0.007 | 0.007 | 0 | 0.07 |
| | Mask | [87.68, 118.0] | [-12%, 18%] | 0.007 | 0.007 | 0 | 0 |
| | MPC | [87.68, 115.3] | [-12%, 15%] | 37.65 | 37.65 | 0 | 0 |
| | Shuffle+MPC | [87.68, 115.3] | [-12%, 15%] | 37.65 | 37.65 | 0 | 0 |
| HLL15 | None | [99.15, 100.2] | [-1%, 0%] | 1.44 | 1.44 | 36.12 | 36.12 |
| | Shuffle | [99.15, 100.2] | [-1%, 0%] | 1.44 | 1.44 | 0 | 36.12 |
| | Rehash | [99.15, 100.2] | [-1%, 0%] | 1.6 | 1.625 | 0 | 36.12 |
| | Mask | [48.41, 464.9] | [-52%, 365%] | 1.207 | 1.207 | 0 | 0 |
| Hashed IDs | None | [100.00, 100.00] | [0%, 0%] | 0.002 | 0.002 | 190.6 | 190.6 |
| | Rehash | [100.00, 100.00] | [0%, 0%] | 0.002 | 0.002 | 0 | 190.6 |

**Supplementary Table 4**. 100 patient query. We measure the accuracy, wait time, and privacy risk of running the query using raw counts, HyperLogLog sketches of various sizes, and sending full Hashed patient IDs.

| Method | Obfuscation | Estimated Count | | Wait (s) | | Risk - Hub | Risk - Hub + Hospital |
|---|---|---|---|---|---|---|---|
| | | Absolute | Relative | Mean | Max | | |
| Count | None | [93.47, 1,973] | [-91%, 97%] | 0 | 0 | 43.84 | 44 |
| | Mask | [93.47, 2,236] | [-91%, 124%] | 0 | 0 | 0 | 0 |
| | MPC | [1,863, 1,973] | [86%, 97%] | 0.099 | 0.099 | 0 | 0 |
| HLL0 | None | [177.6, 22,737] | [-82%, 2174%] | 0 | 0 | 0 | 0 |
| | Shuffle | [177.6, 22,737] | [-82%, 2174%] | 0 | 0 | 0 | 0 |
| | Rehash | [177.6, 22,737] | [-82%, 2174%] | 0 | 0.001 | 0 | 0 |
| | Mask | [177.6, 22,737] | [-82%, 2174%] | 0 | 0 | 0 | 0 |
| | MPC | [177.6, 22,737] | [-82%, 2174%] | 0.385 | 0.385 | 0 | 0 |
| | Shuffle+MPC | [177.6, 22,737] | [-82%, 2174%] | 0.385 | 0.385 | 0 | 0 |
| HLL1 | None | [277.9, 5,118] | [-72%, 412%] | 0 | 0 | 0.04 | 0 |
| | Shuffle | [277.9, 5,118] | [-72%, 412%] | 0 | 0 | 0.03 | 0 |
| | Rehash | [277.9, 5,118] | [-72%, 412%] | 0 | 0.001 | 0 | 0.04 |
| | Mask | [237.9, 5,118] | [-76%, 412%] | 0 | 0 | 0 | 0 |
| | MPC | [277.9, 5,118] | [-72%, 412%] | 0.676 | 0.676 | 0 | 0 |
| | Shuffle+MPC | [277.9, 5,118] | [-72%, 412%] | 0.676 | 0.676 | 0 | 0 |
| HLL4 | None | [594.1, 1,498] | [-41%, 50%] | 0.001 | 0.001 | 0.19 | 0 |
| | Shuffle | [594.1, 1,498] | [-41%, 50%] | 0.001 | 0.001 | 0 | 0 |
| | Rehash | [594.1, 1,498] | [-41%, 50%] | 0.001 | 0.002 | 0 | 0 |
| | Mask | [594.1, 1,498] | [-41%, 50%] | 0.001 | 0.001 | 0 | 0 |
| | MPC | [594.1, 1,498] | [-41%, 50%] | 4.764 | 4.764 | 0 | 0 |
| | Shuffle+MPC | [594.1, 1,498] | [-41%, 50%] | 4.764 | 4.764 | 0 | 0 |
| HLL7 | None | [839.9, 1,180] | [-16%, 18%] | 0.006 | 0.006 | 1.67 | 2 |
| | Shuffle | [839.9, 1,180] | [-16%, 18%] | 0.006 | 0.006 | 0 | 2 |
| | Rehash | [839.9, 1,180] | [-16%, 18%] | 0.006 | 0.007 | 0 | 2 |
| | Mask | [832.6, 1,251] | [-17%, 25%] | 0.006 | 0.006 | 0 | 0 |
| | MPC | [839.9, 1,180] | [-16%, 18%] | 37.61 | 37.61 | 0 | 0 |
| | Shuffle+MPC | [839.9, 1,180] | [-16%, 18%] | 37.61 | 37.61 | 0 | 0 |
| HLL15 | None | [992.4, 1,008] | [-1%, 1%] | 1.44 | 1.44 | 375 | 375 |
| | Shuffle | [992.4, 1,008] | [-1%, 1%] | 1.44 | 1.44 | 0 | 375 |
| | Rehash | [992.4, 1,008] | [-1%, 1%] | 1.601 | 1.621 | 0 | 375 |
| | Mask | [93.02, 2,231] | [-91%, 123%] | 0.267 | 0.267 | 0 | 0 |
| Hashed IDs | None | [1000.0, 1000.0] | [0%, 0%] | 0.002 | 0.002 | 1,919 | 1,919 |
| | Rehash | [1000.0, 1000.0] | [0%, 0%] | 0.002 | 0.002 | 0 | 1,919 |

**Supplementary Table 5**. 1,000 patient query. We measure the accuracy, wait time, and privacy risk of running the query using raw counts, HyperLogLog sketches of various sizes, and sending full Hashed patient IDs.

| Method | Obfuscation | Estimated Count | | Wait (s) | | Risk - Hub | Risk - Hub + Hospital |
|---|---|---|---|---|---|---|---|
| | | Absolute | Relative | Mean | Max | | |
| Count | None | [899.9, 19,470] | [-91%, 95%] | 0 | 0 | 2.65 | 2.65 |
| | Mask | [899.9, 19,477] | [-91%, 95%] | 0 | 0 | 0 | 0 |
| | MPC | [18,886, 19,470] | [89%, 95%] | 0.099 | 0.099 | 0 | 0 |
| HLL0 | None | [1,421, 363,799] | [-86%, 3538%] | 0 | 0 | 0.23 | 0.23 |
| | Shuffle | [1,421, 363,799] | [-86%, 3538%] | 0 | 0 | 0.23 | 0.23 |
| | Rehash | [1,421, 363,799] | [-86%, 3538%] | 0.001 | 0.01 | 0 | 0.23 |
| | Mask | [1,421, 182,085] | [-86%, 1721%] | 0 | 0 | 0 | 0 |
| | MPC | [1,421, 363,799] | [-86%, 3538%] | 0.384 | 0.384 | 0 | 0 |
| | Shuffle+MPC | [1,421, 363,799] | [-86%, 3538%] | 0.384 | 0.384 | 0 | 0 |
| HLL1 | None | [2,802, 56,273] | [-72%, 463%] | 0 | 0 | 0.31 | 0.31 |
| | Shuffle | [2,802, 56,273] | [-72%, 463%] | 0 | 0 | 0.23 | 0.31 |
| | Rehash | [2,802, 56,273] | [-72%, 463%] | 0.001 | 0.01 | 0 | 0.31 |
| | Mask | [2,559, 41,028] | [-74%, 310%] | 0 | 0 | 0 | 0 |
| | MPC | [2,802, 56,273] | [-72%, 463%] | 0.676 | 0.676 | 0 | 0 |
| | Shuffle+MPC | [2,802, 56,273] | [-72%, 463%] | 0.676 | 0.676 | 0 | 0 |
| HLL4 | None | [6,243, 16,850] | [-38%, 68%] | 0.001 | 0.001 | 1.84 | 1.84 |
| | Shuffle | [6,243, 16,850] | [-38%, 68%] | 0.001 | 0.001 | 0.23 | 1.84 |
| | Rehash | [6,243, 16,850] | [-38%, 68%] | 0.002 | 0.011 | 0 | 1.84 |
| | Mask | [5,704, 16,369] | [-43%, 64%] | 0.001 | 0.001 | 0 | 0 |
| | MPC | [6,243, 16,850] | [-38%, 68%] | 4.787 | 4.787 | 0.05 | 0.05 |
| | Shuffle+MPC | [6,243, 16,850] | [-38%, 68%] | 4.787 | 4.787 | 0 | 0.05 |
| HLL7 | None | [8,310, 11,347] | [-17%, 13%] | 0.006 | 0.006 | 15.73 | 15.73 |
| | Shuffle | [8,310, 11,347] | [-17%, 13%] | 0.006 | 0.006 | 0.23 | 15.73 |
| | Rehash | [8,310, 11,347] | [-17%, 13%] | 0.007 | 0.016 | 0 | 15.73 |
| | Mask | [7,167, 14,123] | [-28%, 41%] | 0.005 | 0.005 | 0 | 0 |
| | MPC | [8,310, 11,347] | [-17%, 13%] | 37.83 | 37.83 | 0.3 | 0.3 |
| | Shuffle+MPC | [8,310, 11,347] | [-17%, 13%] | 37.83 | 37.83 | 0 | 0.3 |
| HLL15 | None | [9,928, 10,075] | [-1%, 1%] | 1.462 | 1.462 | 3,707 | 3,707 |
| | Shuffle | [9,928, 10,075] | [-1%, 1%] | 1.462 | 1.462 | 0.23 | 3,707 |
| | Rehash | [9,928, 10,075] | [-1%, 1%] | 1.625 | 1.668 | 0 | 3,707 |
| | Mask | [899.9, 19,477] | [-91%, 95%] | 0.012 | 0.012 | 0 | 0 |
| Hashed IDs | None | [10,000, 10,000] | [0%, 0%] | 0.002 | 0.002 | 19,174 | 19,174 |
| | Rehash | [10,000, 10,000] | [0%, 0%] | 0.002 | 0.004 | 0 | 19,174 |

**Supplementary Table 6**. 10,000 patient query. We measure the accuracy, wait time, and privacy risk of running the query using raw counts, HyperLogLog sketches of various sizes, and sending full Hashed patient IDs.

| Method | Obfuscation | Estimated Count | | Wait (s) | | Risk - Hub | Risk - Hub + Hospital |
|---|---|---|---|---|---|---|---|
| | | Absolute | Relative | Mean | Max | | |
| Count | None | [9,057, 194,126] | [-91%, 94%] | 0 | 0 | 0.02 | 0.02 |
| | Mask | [9,057, 194,126] | [-91%, 94%] | 0 | 0 | 0 | 0 |
| | MPC | [188,850, 194,126] | [89%, 94%] | 0.099 | 0.099 | 0 | 0 |
| HLL0 | None | [11,369, 727,598] | [-89%, 628%] | 0 | 0 | 1.32 | 1.32 |
| | Shuffle | [11,369, 727,598] | [-89%, 628%] | 0 | 0 | 1.32 | 1.32 |
| | Rehash | [11,369, 727,598] | [-89%, 628%] | 0.011 | 0.096 | 0 | 1.32 |
| | Mask | [11,369, 369,142] | [-89%, 269%] | 0 | 0 | 0 | 0 |
| | MPC | [11,369, 727,598] | [-89%, 628%] | 0.385 | 0.385 | 0 | 0 |
| | Shuffle+MPC | [11,369, 727,598] | [-89%, 628%] | 0.385 | 0.385 | 0 | 0 |
| HLL1 | None | [20,470, 393,030] | [-80%, 293%] | 0 | 0 | 2.71 | 2.71 |
| | Shuffle | [20,470, 393,030] | [-80%, 293%] | 0 | 0 | 1.32 | 2.71 |
| | Rehash | [20,470, 393,030] | [-80%, 293%] | 0.011 | 0.115 | 0 | 2.71 |
| | Mask | [20,470, 226,929] | [-80%, 127%] | 0 | 0 | 0 | 0 |
| | MPC | [20,470, 393,030] | [-80%, 293%] | 0.677 | 0.677 | 0 | 0 |
| | Shuffle+MPC | [20,470, 393,030] | [-80%, 293%] | 0.677 | 0.677 | 0 | 0 |
| HLL4 | None | [58,024, 169,675] | [-42%, 70%] | 0.001 | 0.001 | 19.51 | 20 |
| | Shuffle | [58,024, 169,675] | [-42%, 70%] | 0.001 | 0.001 | 1.32 | 19.51 |
| | Rehash | [58,024, 169,675] | [-42%, 70%] | 0.012 | 0.123 | 0 | 19.51 |
| | Mask | [45,325, 138,479] | [-55%, 38%] | 0.001 | 0.001 | 0 | 0 |
| | MPC | [58,024, 169,675] | [-42%, 70%] | 4.771 | 4.771 | 0 | 0 |
| | Shuffle+MPC | [58,024, 169,675] | [-42%, 70%] | 4.771 | 4.771 | 0 | 0 |
| HLL7 | None | [79,547, 117,654] | [-20%, 18%] | 0.006 | 0.006 | 158 | 158 |
| | Shuffle | [79,547, 117,654] | [-20%, 18%] | 0.006 | 0.006 | 1.32 | 158.3 |
| | Rehash | [79,547, 117,654] | [-20%, 18%] | 0.017 | 0.103 | 0 | 158 |
| | Mask | [15,973, 190,297] | [-84%, 90%] | 0.001 | 0.001 | 0 | 0 |
| | MPC | [79,547, 117,654] | [-20%, 18%] | 37.63 | 37.63 | 1.64 | 1.64 |
| | Shuffle+MPC | [79,547, 117,654] | [-20%, 18%] | 37.63 | 37.63 | 0 | 1.64 |
| HLL15 | None | [99,893, 101,691] | [-0%, 2%] | 1.363 | 1.363 | 36,876 | 36,876 |
| | Shuffle | [99,893, 101,691] | [-0%, 2%] | 1.363 | 1.363 | 1 | 36,876 |
| | Rehash | [99,893, 101,691] | [-0%, 2%] | 1.537 | 1.637 | 0 | 36,876 |
| | Mask | [9,057, 194,126] | [-91%, 94%] | 0.001 | 0.001 | 0 | 0 |
| Hashed IDs | None | [100,000, 100,000] | [0%, 0%] | 0.008 | 0.008 | 191,759 | 191,759 |
| | Rehash | [100,000, 100,000] | [0%, 0%] | 0.011 | 0.032 | 0 | 191,759 |

**Supplementary Table 7**. 100,000 patient query. We measure the accuracy, wait time, and privacy risk of running the query using raw counts, HyperLogLog sketches of various sizes, and sending full Hashed patient IDs.

| Method | Obfuscation | Estimated Count | | Wait (s) | | Risk - Hub | Risk - Hub + Hospital |
|---|---|---|---|---|---|---|---|
| | | Absolute | Relative | Mean | Max | | |
| Count | None | [90,222, 1,940,247] | [-91%, 94%] | 0 | 0 | 0 | 0 |
| | Mask | [90,222, 1,940,247] | [-91%, 94%] | 0 | 0 | 0 | 0 |
| | MPC | [1,888,932, 1,940,247] | [89%, 94%] | 0.1 | 0.1 | 0 | 0 |
| HLL0 | None | [181,899, 17,753,385] | [-82%, 1675%] | 0 | 0 | 11.71 | 11.71 |
| | Shuffle | [181,899, 17,753,385] | [-82%, 1675%] | 0 | 0 | 11.71 | 11.71 |
| | Rehash | [181,899, 17,753,385] | [-82%, 1675%] | 0.104 | 0.947 | 0 | 11.71 |
| | Mask | [90,950, 932,550] | [-91%, -7%] | 0 | 0 | 0 | 0 |
| | MPC | [181,899, 17,753,385] | [-82%, 1675%] | 0.385 | 0.385 | 0.2 | 0.2 |
| | Shuffle+MPC | [181,899, 17,753,385] | [-82%, 1675%] | 0.385 | 0.385 | 0.2 | 0.2 |
| HLL1 | None | [206,887, 11,581,296] | [-79%, 1058%] | 0 | 0 | 23.06 | 23.06 |
| | Shuffle | [206,887, 11,581,296] | [-79%, 1058%] | 0 | 0 | 12.15 | 23.06 |
| | Rehash | [206,887, 11,581,296] | [-79%, 1058%] | 0.105 | 0.947 | 0 | 23.06 |
| | Mask | [109,175, 1,273,729] | [-89%, 27%] | 0 | 0 | 0 | 0 |
| | MPC | [206,887, 11,581,296] | [-79%, 1058%] | 0.679 | 0.679 | 0.39 | 0.39 |
| | Shuffle+MPC | [206,887, 11,581,296] | [-79%, 1058%] | 0.679 | 0.679 | 0.2 | 0.39 |
| HLL4 | None | [538,869, 1,713,823] | [-46%, 71%] | 0.001 | 0.001 | 186.3 | 186.3 |
| | Shuffle | [538,869, 1,713,823] | [-46%, 71%] | 0.001 | 0.001 | 12.41 | 186.3 |
| | Rehash | [538,869, 1,713,823] | [-46%, 71%] | 0.105 | 0.943 | 0 | 186.3 |
| | Mask | [90,644, 1,892,176] | [-91%, 89%] | 0 | 0 | 0 | 0 |
| | MPC | [538,869, 1,713,823] | [-46%, 71%] | 4.754 | 4.754 | 1.91 | 1.91 |
| | Shuffle+MPC | [538,869, 1,713,823] | [-46%, 71%] | 4.754 | 4.754 | 0.2 | 1.91 |
| HLL7 | None | [837,269, 1,158,432] | [-16%, 16%] | 0.006 | 0.006 | 1,508 | 1,508 |
| | Shuffle | [837,269, 1,158,432] | [-16%, 16%] | 0.006 | 0.006 | 12.48 | 1,508 |
| | Rehash | [837,269, 1,158,432] | [-16%, 16%] | 0.111 | 0.955 | 0 | 1,508 |
| | Mask | [90,222, 1,940,247] | [-91%, 94%] | 0.001 | 0.001 | 0 | 0 |
| | MPC | [837,269, 1,158,432] | [-16%, 16%] | 37.39 | 37.39 | 14.71 | 14.71 |
| | Shuffle+MPC | [837,269, 1,158,432] | [-16%, 16%] | 37.39 | 37.39 | 0.2 | 14.71 |
| HLL15 | None | [987,401, 1,011,968] | [-1%, 1%] | 1.379 | 1.379 | 348,258 | 348,258 |
| | Shuffle | [987,401, 1,011,968] | [-1%, 1%] | 1.379 | 1.379 | 12.5 | 348,258 |
| | Rehash | [987,401, 1,011,968] | [-1%, 1%] | 1.652 | 2.537 | 0 | 348,258 |
| | Mask | [90,222, 1,940,247] | [-91%, 94%] | 0.001 | 0.001 | 0 | 0 |
| Hashed IDs | None | [1,000,000, 1,000,000] | [0%, 0%] | 0.071 | 0.071 | 1,917,817 | 1,917,817 |
| | Rehash | [1,000,000, 1,000,000] | [0%, 0%] | 0.097 | 0.302 | 0 | 1,917,817 |

**Supplementary Table 8**. 1,000,000 patient query. We measure the accuracy, wait time, and privacy risk of running the query using raw counts, HyperLogLog sketches of various sizes, and sending full Hashed patient IDs.

| Method | Obfuscation | Estimated Count | | Wait (s) | | Risk - Hub | Risk - Hub + Hospital |
|---|---|---|---|---|---|---|---|
| | | Absolute | Relative | Mean | Max | | |
| Count | None | [902,793, 19,402,938] | [-91%, 94%] | 0 | 0 | 0 | 0 |
| | Mask | [902,793, 19,402,938] | [-91%, 94%] | 0 | 0 | 0 | 0 |
| | MPC | [18,884,586, 19,402,938] | [89%, 94%] | 0.105 | 0.105 | 0 | 0 |
| HLL0 | None | [1,455,196, 71,013,541] | [-85%, 610%] | 0 | 0 | 70.06 | 70 |
| | Shuffle | [1,455,196, 71,013,541] | [-85%, 610%] | 0 | 0 | 70 | 70 |
| | Rehash | [1,455,196, 71,013,541] | [-85%, 610%] | 1.032 | 9.685 | 0 | 70 |
| | Mask | [734,987, 17,239,998] | [-93%, 72%] | 0 | 0 | 0 | 0 |
| | MPC | [1,455,196, 71,013,541] | [-85%, 610%] | 0.383 | 0.383 | 1 | 1 |
| | Shuffle+MPC | [1,455,196, 71,013,541] | [-85%, 610%] | 0.383 | 0.383 | 1 | 1 |
| HLL1 | None | [2,215,613, 41,923,242] | [-78%, 319%] | 0 | 0 | 140.8 | 141 |
| | Shuffle | [2,215,613, 41,923,242] | [-78%, 319%] | 0 | 0 | 92.64 | 140.8 |
| | Rehash | [2,215,613, 41,923,242] | [-78%, 319%] | 1.033 | 9.483 | 0 | 141 |
| | Mask | [887,447, 18,931,827] | [-91%, 89%] | 0 | 0 | 0 | 0 |
| | MPC | [2,215,613, 41,923,242] | [-78%, 319%] | 0.675 | 0.675 | 2 | 2 |
| | Shuffle+MPC | [2,215,613, 41,923,242] | [-78%, 319%] | 0.675 | 0.675 | 1.17 | 2 |
| HLL4 | None | [6,171,672, 16,428,565] | [-38%, 64%] | 0.001 | 0.001 | 1,123 | 1,123 |
| | Shuffle | [6,171,672, 16,428,565] | [-38%, 64%] | 0.001 | 0.001 | 122 | 1,123 |
| | Rehash | [6,171,672, 16,428,565] | [-38%, 64%] | 1.035 | 9.495 | 0 | 1,123 |
| | Mask | [902,793, 19,402,938] | [-91%, 94%] | 0.001 | 0.001 | 0 | 0 |
| | MPC | [6,171,672, 16,428,565] | [-38%, 64%] | 4.755 | 4.755 | 18 | 18 |
| | Shuffle+MPC | [6,171,672, 16,428,565] | [-38%, 64%] | 4.755 | 4.755 | 1.18 | 18 |
| HLL7 | None | [8,352,290, 12,307,282] | [-16%, 23%] | 0.006 | 0.006 | 8,947 | 8,947 |
| | Shuffle | [8,352,290, 12,307,282] | [-16%, 23%] | 0.006 | 0.006 | 126 | 8,947 |
| | Rehash | [8,352,290, 12,307,282] | [-16%, 23%] | 1.043 | 9.496 | 0 | 8,947 |
| | Mask | [902,793, 19,402,938] | [-91%, 94%] | 0.001 | 0.001 | 0 | 0 |
| | MPC | [8,352,290, 12,307,282] | [-16%, 23%] | 37.41 | 37.41 | 144 | 144 |
| | Shuffle+MPC | [8,352,290, 12,307,282] | [-16%, 23%] | 37.41 | 37.41 | 1.18 | 144 |
| HLL15 | None | [9,887,923, 10,123,720] | [-1%, 1%] | 1.392 | 1.392 | 2,127,583 | 2,127,583 |
| | Shuffle | [9,887,923, 10,123,720] | [-1%, 1%] | 1.392 | 1.392 | 127 | 2,127,583 |
| | Rehash | [9,887,923, 10,123,720] | [-1%, 1%] | 2.641 | 11.55 | 0 | 2,127,583 |
| | Mask | [902,793, 19,402,938] | [-91%, 94%] | 0.001 | 0.001 | 0 | 0 |
| Hashed IDs | None | [10,000,000, 10,000,000] | [0%, 0%] | 0.794 | 0.794 | 19,178,114 | 19,178,114 |
| | Rehash | [10,000,000, 10,000,000] | [0%, 0%] | 1.045 | 3.09 | 0 | 19,178,114 |

**Supplementary Table 9**. 10,000,000 patient query. We measure the accuracy, wait time, and privacy risk of running the query using raw counts, HyperLogLog sketches of various sizes, and sending full Hashed patient IDs.

| Method | Obfuscation | Estimated Count | | Wait (s) | | Risk - Hub | Risk - Hub + Hospital |
|---|---|---|---|---|---|---|---|
| | | Absolute | Relative | Mean | Max | | |
| Count | None | [9,431,902, 200,018,431] | [-91%, 100%] | 0 | 0 | 0 | 0 |
| | Mask | [9,431,902, 200,018,431] | [-91%, 100%] | 0 | 0 | 0 | 0 |
| | MPC | [199,984,637, 200,018,431] | [100%, 100%] | 0.171 | 0.171 | 0 | 0 |
| HLL0 | None | [11,641,564, 818,287,824] | [-88%, 718%] | 0 | 0 | 99.97 | 99.97 |
| | Shuffle | [11,641,564, 818,287,824] | [-88%, 718%] | 0 | 0 | 99.97 | 99.97 |
| | Rehash | [11,641,564, 818,287,824] | [-88%, 718%] | 9.42 | 84.36 | 0 | 99.97 |
| | Mask | [9,431,902, 200,018,431] | [-91%, 100%] | 0.001 | 0.001 | 0 | 0 |
| | MPC | [11,641,564, 818,287,824] | [-88%, 718%] | 0.387 | 0.387 | 9.91 | 9.91 |
| | Shuffle+MPC | [11,641,564, 818,287,824] | [-88%, 718%] | 0.387 | 0.387 | 9.91 | 9.91 |
| HLL1 | None | [22,952,975, 448,641,535] | [-77%, 349%] | 0 | 0 | 199.9 | 199.9 |
| | Shuffle | [22,952,975, 448,641,535] | [-77%, 349%] | 0 | 0 | 198.5 | 199.9 |
| | Rehash | [22,952,975, 448,641,535] | [-77%, 349%] | 9.42 | 84.36 | 0 | 199.9 |
| | Mask | [9,431,902, 200,018,431] | [-91%, 100%] | 0.001 | 0.001 | 0 | 0 |
| | MPC | [22,952,975, 448,641,535] | [-77%, 349%] | 0.681 | 0.681 | 19.82 | 19.82 |
| | Shuffle+MPC | [22,952,975, 448,641,535] | [-77%, 349%] | 0.681 | 0.681 | 10.75 | 19.82 |
| HLL4 | None | [57,717,266, 167,663,627] | [-42%, 68%] | 0.001 | 0.001 | 1,599 | 1,599 |
| | Shuffle | [57,717,266, 167,663,627] | [-42%, 68%] | 0.001 | 0.001 | 854.1 | 1,599 |
| | Rehash | [57,717,266, 167,663,627] | [-42%, 68%] | 9.422 | 84.36 | 0 | 1,599 |
| | Mask | [9,431,902, 200,018,431] | [-91%, 100%] | 0.001 | 0.001 | 0 | 0 |
| | MPC | [57,717,266, 167,663,627] | [-42%, 68%] | 4.768 | 4.768 | 153.8 | 153.8 |
| | Shuffle+MPC | [57,717,266, 167,663,627] | [-42%, 68%] | 4.768 | 4.768 | 11.69 | 153.8 |
| HLL7 | None | [84,484,568, 117,231,365] | [-16%, 17%] | 0.006 | 0.006 | 12,795 | 12,795 |
| | Shuffle | [84,484,568, 117,231,365] | [-16%, 17%] | 0.006 | 0.006 | 1,199 | 12,795 |
| | Rehash | [84,484,568, 117,231,365] | [-16%, 17%] | 9.426 | 84.36 | 0 | 12,795 |
| | Mask | [9,431,902, 200,018,431] | [-91%, 100%] | 0.001 | 0.001 | 0 | 0 |
| | MPC | [84,484,568, 117,231,365] | [-16%, 17%] | 37.43 | 37.43 | 1,246 | 1,246 |
| | Shuffle+MPC | [84,484,568, 117,231,365] | [-16%, 17%] | 37.43 | 37.43 | 11.83 | 1,246 |
| HLL15 | None | [99,097,598, 101,037,154] | [-1%, 1%] | 1.362 | 1.362 | 3,259,256 | 3,259,256 |
| | Shuffle | [99,097,598, 101,037,154] | [-1%, 1%] | 1.362 | 1.362 | 1,265 | 3,259,256 |
| | Rehash | [99,097,598, 101,037,154] | [-1%, 1%] | 10.96 | 85.89 | 0 | 3,259,256 |
| | Mask | [9,431,902, 200,018,431] | [-91%, 100%] | 0.001 | 0.001 | 0 | 0 |
| Hashed IDs | None | [100,000,000, 100,000,000] | [0%, 0%] | 10.12 | 10.12 | 199,999,801 | 199,999,801 |
| | Rehash | [100,000,000, 100,000,000] | [0%, 0%] | 11.56 | 23.31 | 0 | 199,999,801 |

**Supplementary Table 10**. 100,000,000 patient query. We measure the accuracy, wait time, and privacy risk of running the query using raw counts, HyperLogLog sketches of various sizes, and sending full Hashed patient IDs.
